# Sex-specific control of feeding and defensive behaviors by MC3R neurons in the bed nuclei of the stria terminalis

**DOI:** 10.64898/2026.02.08.702109

**Authors:** Michelle N. Bedenbaugh, Marie A. Doyle, Caitlyn M. Edwards, Nick Petersen, Dollada Srisai, Sydney H. Hawkins, Nicholas T. Low, Haley Mendoza-Romero, Jordan A. Brown, Alexander S. Vardy, Danny G. Winder, Richard B. Simerly

## Abstract

The bed nuclei of the stria terminalis (BST) is a nuclear complex that coordinates neuroendocrine, autonomic, and behavioral responses associated with maintaining homeostasis. Here, we demonstrate that melanocortin 3 receptor (MC3R) neurons in the BST (BST^MC3R^) play a key role in modulating feeding behaviors and responses to stressful events. BST^MC3R^ neurons are primarily GABAergic and colocalize with neuropeptides known to regulate feeding and affective behaviors. Chemogenetic activation of BST^MC3R^ neurons causes reduced feeding, particularly in males. BST^MC3R^ neurons are also robustly activated by stress and can control responses to stress and defensive behaviors in both sexes. Whole-brain monosynaptic rabies tracing identified multiple sexually dimorphic inputs to BST^MC3R^ neurons that may contribute to observed functional sexual dimorphisms. Altogether, these data reveal that BST^MC3R^ neurons participate in sexually dimorphic circuits that influence feeding and defensive behaviors, responses to stress, and may represent a potential therapeutic target for stress- and eating-related disorders.

## INTRODUCTION

The BST is a heterogeneous, highly differentiated nuclear complex whereby autonomic, emotional and neuroendocrine signals are integrated and subsequently relayed to regulate the expression of motivated behaviors, including feeding and defensive actions^1–6^. The BST encompasses between 12-18 different nuclei and includes regions both dorsal and ventral to the anterior commissure^7,8^. Dorsal regions of the anteromedial, anterolateral, oval and juxtacapsular nuclei have been implicated in the control of food intake, energy balance homeostasis, ingestive behaviors, responses to stress and anxiety-like behaviors^2,3,9^. The BST receives inputs from many brain regions, including cortical areas such as the prefrontal region, insula and hippocampal formation, as well as inputs from multiple amygdalar nuclei^10^. These autonomic, emotional and neuroendocrine signals are integrated and subsequently relayed to several hypothalamic, thalamic, midbrain and brainstem nuclei to influence behavioral state^1–4,11^. Of interest, the BST is centrally involved in stress-related psychopathologies, such as pathological and adaptive anxiety^5,12–17^. This area has also been implicated in the regulation of food intake and energy balance^3,18,19^ with both anorexic^20–22^ and binge-like eating effects^23^ reported. However, the neural substrates within the BST that may be integrating information driving feeding, stress and anxiety-like behaviors are not well defined.

The central melanocortin system is fundamentally important for controlling energy balance and maintaining homeostasis. Key components of this system are endogenous agonists derived from the prohormone, proopiomelanocortin (POMC), the endogenous antagonist, agouti-related protein (AgRP) and two receptors: MC3R and melanocortin 4 receptor (MC4R)^24,25^. While there is abundant evidence that MC4R plays a determinant role in the central regulation of food intake and energy homeostasis^24,26–30^, the homeostatic functions of MC3R are just beginning to be characterized. MC3R has been implicated in the bidirectional control of responses to homeostatic challenges, providing rheostatic control of energy storage^31^. Importantly, it was also recently discovered that MC3R neurons bidirectionally regulate feeding and anxiety-like behaviors, and global deletion of MC3R produces multiple forms of sexually dimorphic anxiety-related hypophagia^32^. MC3R neurons and terminals are abundantly expressed in the BST^33^, and while MC4R is expressed in the BST, BST^MC4R^ neurons do not appear to be a downstream target for AgRP-driven hunger^34^. Therefore, we hypothesized that the BST^MC3R^ circuitry may function as a sexually dimorphic integration hub that receives a variety of convergent inputs required for effective coordination of feeding, stress and anxiety-like behaviors.

To test this hypothesis, we combined cell-type-specific manipulations with real-time activity monitoring and comprehensive circuit mapping to characterize MC3R neurons in the dorsal BST (BSTd) and define their role in coordinating feeding, stress, and anxiety-like behaviors. Our findings reveal that BST^MC3R^ neurons are activated during, and can influence, feeding, stress and defensive behaviors, but appear to function differently in males and females. Finally, BST^MC3R^ neurons share strong bidirectional connections with brain regions that induce feeding and affective behaviors. However, only the inputs to BST^MC3R^ neurons are sexually dimorphic. These results provide new insights into the multiple functional roles for BST^MC3R^ neurons and identify potential targets for treating eating and stress-related disorders.

## MATERIALS AND METHODS

### Animals

Transgenic BAC mice expressing Cre recombinase under control of the *Mc3r* promoter (MC3R-Cre mice) are maintained in our colony at Vanderbilt University^31^. Mice expressing the Cre-dependent fluorescent reporter tdTomato (tdTom mice; Ai14D-Gt(Rosa)26Sor; stock number: 007914) were obtained from The Jackson Laboratory (Bar Harbor, ME). Wild-type C57BL/6J mice (stock number: 000664) were also obtained from The Jackson Laboratory. To visualize neurons that express MC3R, MC3R-Cre mice were crossed with tdTom mice to generate MC3R-Cre::tdTom mice.

All animal care and experimental procedures were performed in accordance with the guidelines of the National Institutes of Health and the Institutional Care and Use Committee of Vanderbilt University. Mice were housed at 22°C on a 12:12 h light:dark cycle (lights on at 6:00 AM: lights off at 6:00 PM). Mice were provided *ad libitum* access to a standard chow diet (PicoLab Rodent Diet 20 #5053). Mice were weaned at P22 and maintained with mixed genotype littermates until used for experiments.

### Tissue Processing

Unless otherwise stated, mice were anesthetized with tribromoethanol and perfused transcardially with saline followed by fixative (4% paraformaldehyde in borate buffer, pH 9.5). Brains were post-fixed in a solution of 20% sucrose in fixative and cryoprotected in 20% sucrose in 0.2M potassium phosphate buffered saline (KPBS). Four series of 30 µm-thick frozen sections were collected using a sliding microtome and stored in cryoprotectant until used. For DREADD and fiber photometry behavioral cohorts, the presence of fluorescent transgenes and optical fiber placements (fiber photometry) was confirmed before inclusion in the presented datasets.

### Viral Vectors

Adeno-associated viral vectors used in these experiments included AAVDJ-hSyn-FLEX-mGFP-2A-Synaptophysin-mRuby (Stanford Gene Vector and Virus Core, Palo Alto, CA), AAV1-syn-FLEX-splitTVA-EGFP-tTa (Addgene, Watertown, MA), AAV1-TREtight-mTagBFP2-B19G (Addgene), EnvA-G-DeletedRabies-mCherry (Salk Gene Transfer, Targeting and Therapeutics Viral Vector Core, La Jolla, CA), AAV5-hSyn-DIO-hM3D(Gq)-mCherry (Addgene), AAV5-hSyn-DIO-mCherry (Addgene), and AAV5-Syn-FLEX-GCaMP7f (Addgene). AAVs were used as received except for AAV1-syn-FLEX-splitTVA-EGFP-tTa and AAV1-TREtight-mTagBFP2-B19G which were diluted 1:200 and 1:20, respectively in sterile, filtered saline and subsequently mixed 50/50 by volume.

### Viral Injections and Stereotaxic Surgery

Mice (>8 weeks) were anesthetized with isoflurane (5% initial dose, 2% maintenance dose) for intracranial recombinant AAV injection surgeries using a Kopf mouse stereotax model 1900 (David Kopf Instruments, Tujunga, CA). A micro-precision drill was used to drill a small burr-hole directly above the viral injection site. All injections were unilateral and used the same coordinates to target the dorsal region of the BST on the right side (A/P: 0.43mm from bregma, M/L: 0.55mm and D/V: −3.5mm from surface of brain). Mice were injected with 200nL of the indicated AAV at a rate of 50nL/min driven by a Micro4 MicroSyringe pump (World Precision Instruments, Sarasota, FL) into the BST. The needle remained in place for an additional 5 min after injection to prevent viral spread outside of the BST. Mice used in DREADD and fiber photometry experiments recovered for at least 2 weeks before experiments were conducted. For anterograde tracing experiments, tissue from mice that received AAVDJ-hSyn-FLEX-mGFP-2A-Synaptophysin-mRuby was collected six weeks after injection. For monosynaptic rabies retrograde tracing experiments, a 50/50 mixture of AAV1-syn-FLEX-splitTVA-EGFP-tTa and AAV1-TREtight-mTagBFP2-B19G was injected followed by an injection of EnvA-G-DeletedRabies-mCherry 7 days later. Tissue was collected 7 days after EnvA-G-DeletedRabies-mCherry injection.

#### Fiberoptic Cannula Implantation Procedures

For fiber photometry experiments, a mono fiberoptic cannula (Doric Lenses, Quebec, Canada) was implanted immediately following injection of AAV9-hSyn-FLEX-GCaMP7f. To solidify and stabilize the fiberoptic cannula, the skull was etched with gel etchant (Kerr Dental, Brea, CA) and one carbon steel, nickel-plated micro screw (The Phillips Screw Company, Amesbury MA) was placed in the contralateral parietal plate posterior to the implant hole. The implant was bonded by applying Optibond primer (Patterson Dental, Saint Paul, MN) followed by Optibond adhesive (Patterson Dental). UV light was used to cure the Optibond adhesive and Herculite Unidose enamel (Herculite Products Inc., Emigsville, PA) was molded around the screw, fiberoptic implant and exposed skull. The enamel was subsequently cured with UV light.

### Immunohistochemistry, Image Acquisition and Analysis

#### MC3R neuronal distribution

To visualize MC3R neuronal labeling, brains from adult MC3R-Cre::tdTomato mice were collected and processed for immunofluorescence. Sections throughout the rostrocaudal extent of the BST in each animal were rinsed in KPBS and blocked in 2% normal goat serum containing 0.3% Tritonx-100 overnight at 4°C. Sections were then incubated in primary antibody (Rabbit anti-RFP, 1:10,000; Rockland Immunochemicals Inc., Limerick, PA) for 48 h at 4°C. Following primary antibody incubation, sections were rinsed several times in KPBS and incubated in Alexa 568 Goat anti-rabbit secondary antibody (1:500; Life Technologies, Carlsbad, CA) for 1 h at room temperature. Sections were rinsed several times in KPBS and incubated in NeuroTrace 500/524 Green fluorescent Nissl stain (1:500; ThermoFisher Scientific, Waltham, MA) for 20 min at room temperature to aid in visualization of brain cytoarchitecture. Sections were rinsed several times in KPBS, mounted on gelatin-subbed slides and coverslipped using ProLong mounting medium (Life Technologies).

Images from each animal were obtained using a laser scanning confocal microscope (Zeiss LSM 800, Zeiss, Oberkochen, Germany). Image stacks captured to compare neuronal labeling, density and distribution were collected through the z-axis at a frequency of 3.41 µm using a 10x objective (NA 0.45). For a quantitative comparison of cell densities in male and female mice, confocal image stacks were collected through the z-axis at a frequency of 0.8 µm using a 20x objective (NA 0.8). Cytoarchitectonic features of each specific anatomical area analyzed, visualized with the NeuroTrace fluorescent Nissl stain, were used to define matching regions of interest (ROI), and three-dimensional representations of labeled cells from matched sections were digitally rendered using Imaris software (version 9.5.1, Bitplane, Zurich, Switzerland). The total number of MC3R neurons in each area was quantified using the spots function in Imaris.

### RNAscope fluorescence *in situ* hybridization

Four series of 20 µm-thick frozen sections from male and female Wild-type C57BL/6J mice were collected using a sliding microtome. Sections throughout the rostrocaudal extent of the BST in each animal were mounted onto SuperFrost Plus slides (Fisher Scientific), and *in situ* hybridization was performed according to the RNAscope multiplex fluorescent V2 kit user manual for fixed frozen tissue (Advanced Cell Diagnostics, Newark, CA) using RNAscope Probe-Mm-*Mc3r*-C1(Cat# 412541), Probe-Mm-*Slc32a1*-C3(Cat# 319191), Probe-Mm-*Slc17a6*-C2(Cat# 319171), Probe-Mm-*Crh*-C2(Cat# 316091), Probe-Mm-*Prkcd*-C3(Cat# 441791), Probe-Mm-*Th*-C2(Cat# 317621), Probe-Mm-*Glp1r*-C3(Cat# 418851), and Probe-Mm-*Esr1*-C2(Cat# 478201). Slides were coverslipped using ProLong mounting medium (Life Technologies). Images of the dorsal BST (BSTd) of each animal were obtained using a laser scanning confocal microscope (Zeiss LSM 800). Confocal image stacks were collected through the z-axis at a frequency of 0.41 µm using a 40x objective (NA 1.40). Three dimensional representations of labeled cells were digitally rendered using Imaris software (version 9.5.1, Bitplane). To determine overall *Mc3r* mRNA abundance in the BSTd, a region of interest (ROI) was placed around the area, and the total amount of *Mc3r* was quantified using the spots function. Total numbers of labeled, *Slc32a1*, *Slc17a6*, *Crh*, *Prkcd*, *Glp1r*, *Th*, and *Esr1* neurons that co-express *Mc3r* were counted manually in each BSTd image stack, aided by Imaris software (Bitplane, v9.5.1). Only neurons with three times as many fluorescent spots as were measured in background regions, were considered positively labeled for *Mc3r* mRNA. Background for each section was determined by placing ten cell-sized ROIs in user-defined areas, where *Mc3r* labeling appeared to be lacking, and averaging the number of spots counted in each background ROI.

### Chemogenetic Experiments and Behaviors

For all chemogenetic behavioral experiments, mice were moved from the vivarium to the behavioral room and allowed to acclimate for at least 2 hours prior to the start of any assay. Behavior was recorded using EthoVision XT video tracking software (Noldus, Wageningen, the Netherlands) equipped with two cameras. Thirty minutes before the start of any experiment, mice were given an i.p. injection of either saline or Clozapine-N-Oxide (CNO, 2mg/kg; Enzo Life Sciences Inc., Farmingdale, NY). Stress and anxiety behavioral assays were completed in this order: novelty suppressed feeding test, open field test, elevated plus maze and restraint stress. At least two days separated each behavioral experiment.

#### Feeding Behavior in a Fed State

For all food experiments, mice were acclimated to feeding using food jars for at least one week before the start of experiments. Mice and food were weighed daily to establish baseline metrics. On the day of the experiment, mice were weighed and food was removed 2 hours before dark onset to standardize the start of feeding for all animals. At dark onset, food was returned and intake measured every hour for 4 hours.

#### Refeeding After a Fast

Similar procedures as described above were used with the exception that mice were fasted overnight prior to experiments and food intake was measured every hour for 3 hours.

#### Novelty Suppressed Feeding Test

Mice were fasted 24 hours prior to testing. NSFT was performed in the same open field arena that was used for OFT (30cm x 30cm). A single pellet of normal chow (3-4 grams) was placed in the center of the open field. All mice were placed in the same corner of the open field and their activity scored using EthoVison XT. Feeding was scored if the mouse took more than one consecutive bite of the food pellet. Animals were determined to be aphagic if no eating occurred during the 10 minutes of testing.

#### Open Field Test

At the start of the test, mice were placed in the same corner of a 30cm x 30cm open field chamber. The center of the arena was marked as the 5cm x 5cm region in the center of the apparatus. Behavior was recorded for 10 minutes using EthoVision XT.

#### Elevated Plus Maze

To perform the elevated plus maze test, mice were initially placed in the closed arm of the arena. The arena consisted of 2, 50cm long arms. One arm was enclosed by walls while the other arm was completely open to surroundings. Activity in the maze was recorded for 10 minutes using EthoVision XT.

#### Restraint Stress

Mice were placed in a RESTRAINT device^35^ for 30 minutes. Struggling behavior was recorded using EthoVision XT and manually scored by a blinded observer. The amount of food eaten in 24 hours after restraint stress was weighed and recorded.

### Fiber Photometry Experiments and Behaviors

Fiber photometry experiments were performed with a Tucker-Davis Technologies RZ5P system (Tucker-Davis Technologies Inc., Alachua, FL) and Synapse software (Tucker-Davis Technologies) as previously described^36–38^. Briefly, excitation light was generated by a 470nm blue light LED and 405nm violet light LED relayed through a filtered fluorescence minicube at spectral bandwidths of 460-495nm and 405nm. Light was delivered to the tissue through a low autofluorescence 0.57NA mono fiberoptic patch cord with a 400µm diameter core (Doric Lenses) secured to the animals’ cannula. Fluorescence emission was collected through the fiber using a femtowatt photoreceiver. The 470nm and 405nm fiber-coupled LEDs were modulated at 210 Hz and 530 Hz, respectively. Light power output of each LED was measured prior to each recording at the fiber tip and maintained at 20µW (405nm LED) and 30µW (470nm LED) by slightly adjusting the level (405nm – 25.7mA ± 1.2 SD; 470nm – 7.7mA ± 0.6 SD) with constant DC offset (405nm – 10mA; 470nm – 4mA) prior to recording each session. All signals were acquired at 1kHz with a 6Hz lowpass.

MATLAB scripts from TDT were used to extract the 405nm and 470nm signals and custom MATLAB scripts were used for further analysis. The 405nm channel was used as an isosbestic, GCaMP-independent control wavelength to correct for photobleaching, movement artifacts, and expression as compared to the GCaMP-dependent 470 nm channel. For each recording, a normalized signal was generated by dividing the 470nm GCaMP-dependent signal by the 405nm GCaMP-independent signal. A ΔF/F trace was then calculated for each recording with the equation:

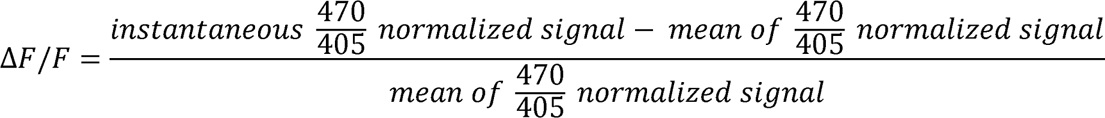

A z-score trace was calculated with the equation:

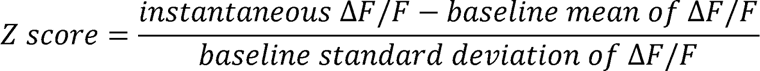

#### Behavioral Experiments

Prior to any experiments being performed, mice were acclimated to attachment of fiberoptic cables for 20 minutes per day in a clean, empty cage for three consecutive days. On test day, mice were transported to the behavioral room and allowed to habituate for one hour before beginning the experiment. The patch cable was connected to the fiber optic implants and animals were habituated in a clean, empty cage for 5 minutes before the start of any behavior test. All behavioral tests were hand scored with the assistance of open-source behavioral software, BORIS^39^. Fiber photometry data were analyzed using custom Python scripts. Calcium transients were aligned to the initiation of a behavioral bout. Activity was analyzed from −3 to 10 seconds.

#### Feeding Behavior in a Fed State

Mice were connected to the TDT system and a baseline recording was captured for 5 minutes. The mice were then exposed to a normal chow pellet placed in the middle of the cage for 5 minutes followed by exposure to a 45% high fat diet pellet (Research Diets D12451) for 5 minutes. Mice were scored as “sniffing” a food pellet if they made contact with the pellet but did not take a bite. Mice were scored as “biting” a food pellet if they took at least two consecutive bites of a food pellet. Each animal’s response to the experimenter’s hand over the cage (looming), as well as their response to rearing up on the side of the cage (rearing) was also recorded.

#### Feeding Behavior in a Fasted State

The same procedures as described above were used with the exception that mice were fasted overnight before conducting experiments.

#### Novel Object

Mice were connected to the TDT system and a baseline recording was captured for 5 minutes. The mice were then exposed to a novel object (battery) placed in the middle of the cage for 5 minutes.

#### Shaker Stress

A custom oscillating orbital shaker with a 2cm orbit was built using 3D printed parts and commercially available hardware and electronic components. The shaker was controlled by an Arduino MEGA microcontroller and consisted of a stepper motor and digital inputs/outputs to relay TTLs to and from the TDT photometry system. Mice were connected to the TDT system and a baseline recording was captured for 5 minutes. The shaker stress protocol was started after the 5-minute baseline period ended through a TTL sent from the TDT system to the Arduino. The shaker stress paradigm consisted of three automated 30 sec successive shaking events of increasing intensities (low speed – 60RPM, medium speed – 120RPM, and high speed – 200RPM) with a 3-minute rest interval. All shaker stress recordings were performed during the light cycle.

### Anterograde Tracing, Image Acquisition and Analysis

To visualize projections of BST^MC3R^ neurons, brains from adult MC3R-Cre mice injected with AAVDJ-hSyn-FLEX-mGFP-2A-Synaptophysin-mRuby were collected and processed for immunofluorescence. One complete series of sections throughout the rostrocaudal extent of the brain in each animal were rinsed in KPBS and blocked in 2% normal goat serum containing 0.3% Tritonx-100 overnight at 4°C. Sections were then incubated in primary antibodies (Rabbit anti-RFP, 1:1,000; Rockland Immunochemicals and Chicken anti-GFP, 1:10,000; Aves Labs Inc., Davis, CA) for 48 h at 4°C. Following primary antibody incubation, sections were rinsed several times in KPBS and incubated in Alexa 568 Goat anti-rabbit secondary antibody and Alexa 488 Goat anti-chicken secondary antibdoy (1:500; Life Technologies) for 1 h at room temperature. Sections were rinsed several times in KPBS and incubated in NeuroTrace 640/660 Deep Red fluorescent Nissl stain (1:200; ThermoFisher Scientific) for 20 min at room temperature to aid in visualization of brain cytoarchitecture. Sections were rinsed several times in KPBS, mounted on gelatin-subbed slides and coverslipped using ProLong mounting medium (Life Technologies).

Images from each animal were obtained using a laser scanning confocal microscope (Zeiss LSM 800). Image stacks captured to compare the number of BST^MC3R^ neurons transfected with virus were collected through the z-axis at a frequency of 3.41 µm using a 10x objective (NA 0.45). For quantitative comparison of terminal densities in male and female mice, confocal image stacks were collected through the z-axis at a frequency of 0.8 µm using a 20x objective (NA 0.8). Cytoarchitectonic features of each specific anatomical area analyzed, visualized with the NeuroTrace fluorescent Nissl stain, were used to define matching regions of interest (ROI), and three-dimensional representations of labeled terminals from matched sections were digitally rendered using Imaris software. The total number of MC3R terminals in each area was quantified using the spots function in Imaris. Terminal density was normalized using the number of BST^MC3R^ neurons that were transfected with virus in each animal.

### Tissue Clearing and Light-sheet Microscopy

#### Tissue Collection, Preparation and Clearing

To spatially map monosynaptic inputs to BST^MC3R^ neurons in intact tissue, brains were stabilized and cleared using the stabilization under harsh conditions via intramolecular epoxide linkages to prevent degradation (SHIELD) protocol ^40^. After tissue collection, brains were incubated in fixative overnight and then transferred to SHIELD-OFF solution (LifeCanvas Technologies, Cambridge, MA) and incubated at 4°C for 3 days. Brains were next incubated in SHIELD-ON solution (LifeCanvas Technologies) for 24 h at 37°C. SHIELD-fixed brains were passively cleared for one week at 42°C in delipidation buffer (LifeCanvas Technologies). Once samples were delipidated, tissue was incubated in EasyIndex (LifeCanvas Technologies) to render the sample optically transparent for imaging.

#### Imaging and Data Processing

SHIELD brains were imaged using an axially swept light-sheet microscope (SmartSPIM, LifeCanvas Technologies) equipped with a 3.6x objective (NA 0.2, uniform axial resolution ∼4 µm) and sCMOS camera with a rolling shutter ^41,42^. Following acquisition, images were stitched to generate composite TIFF images using a modified version of Terastitcher ^43^. Stitched TIFF images were converted to Imaris files using Imaris File Converter 9.2.1, and 3D renderings of brains were visualized using Imaris software.

### Brain Registration and Volumetric Quantification of Monosynaptic Inputs to BST^MC3R^ Neurons

The distribution of neurons in SHIELD cleared brains was registered to the Allen Common Coordinate Reference Framework (ACCF, V3) and regional densities of labeled cells quantified through the application of machine learning filters developed from a user-classified training set using NeUroGlancer Ground Truth (NUGGT), with custom modifications provided by LifeCanvas Technologies ^44^. A minimum of 200 cells per brain were trained using the NUGGT pipeline. The density of cells in each brain region was divided by the total number of cells registered for each brain to normalize for differences in viral transfection and expression.

### Experimental Design and Statistical Analyses

Group data are presented as mean values ± SEM. Statistical significance was determined using GraphPad Prism software (GraphPad Software, San Diego, CA). Statistical parameters, including the number of animals analyzed and the specific statistical tests performed can be found in the figure legends. P-values <0.05 were considered statistically significant.

## RESULTS

### MC3R Neurons are Expressed in the BST and Colocalize with Substrates that Drive Feeding and Affective Behaviors

MC3R neurons are found throughout the rostrocaudal extent of the BST in several distinct subnuclei (Figure 1a-c). In the anterior BST, discrete clusters of MC3R neurons are present in the anteromedial (BSTam; Figure 1a,b), anterolateral (BSTal; Figure 1a,b), oval (BSTov; Figure 1a,b) and juxtacapsular nucleus (BSTju; Figure 1a,b) while little to no MC3R neurons are observed in the fusiform nucleus (BSTfu; Figure 1b). A high density of MC3R neurons is found in the posterior division of the BST, particularly within the principal nucleus (BSTpr; Figure 1c). Because previous literature has identified the BSTam, al, ov and ju as regions heavily involved in the control of food intake, energy balance homeostasis, ingestive behaviors, responses to stress and anxiety-like behaviors ^2,3,9^, the dorsal region of these subnuclei was targeted and analyzed in experiments reported here. For simplicity, the term dorsal BST (BSTd) will be used to describe the BSTam, al, ov and ju that reside dorsal to the anterior commissure.

**Figure 1.**
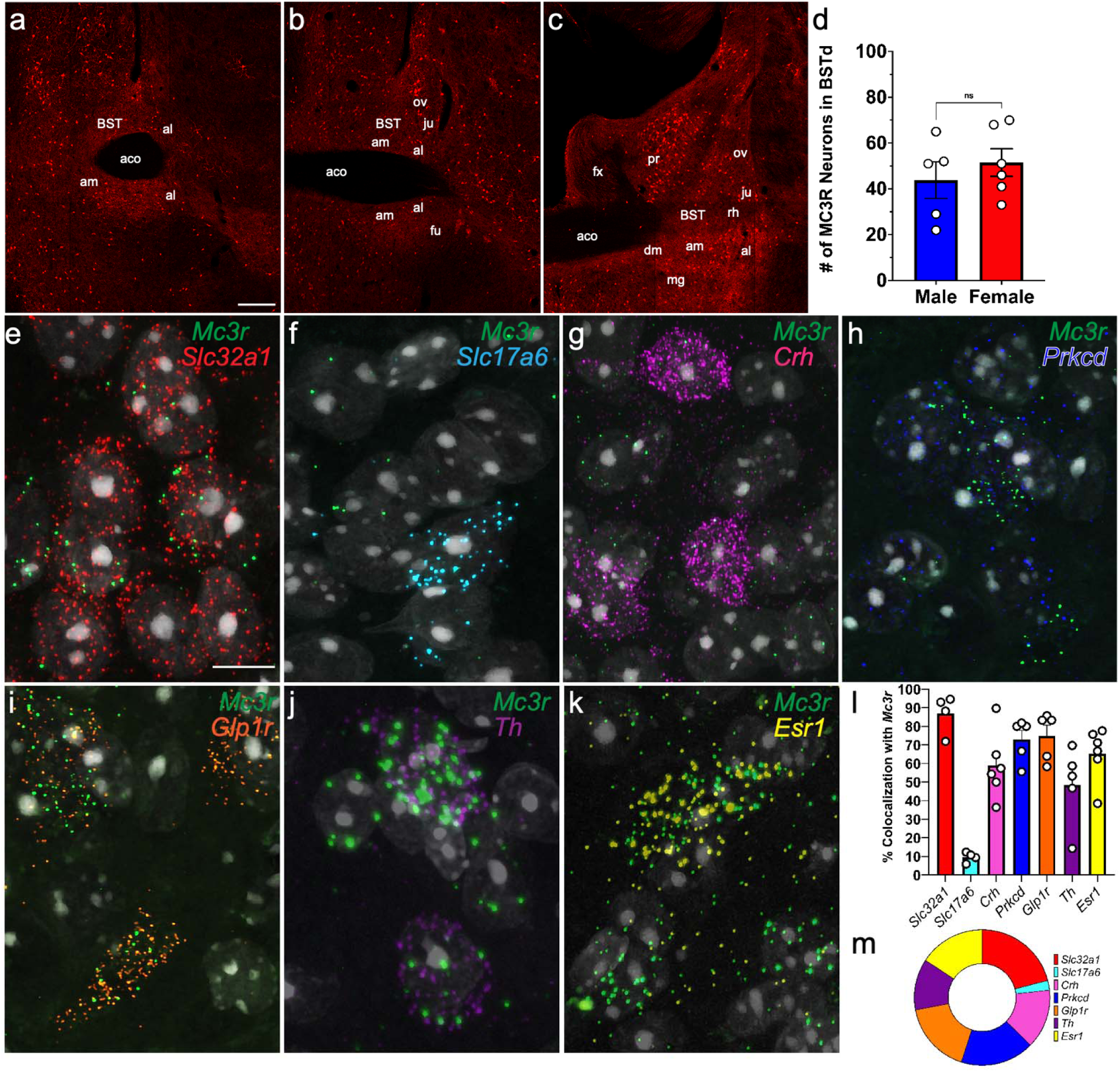
MC3R neurons are found in the BST and coexpress substrates that drive feeding, stress and anxiety-like behaviors. **a, b, c** Low magnification confocal images in an MC3R-Cre::tdTomato mouse illustrating MC3R neurons are distributed in distinct patterns throughout the rostrocaudal extent of the BST. Scale bar, 100 µm. **d** No differences in the number of MC3R neurons was found in the BSTd between males and females (n=5-6/group, unpaired t-test). **e-k** High magnification confocal images depicting *Mc3r* neurons coexpress **e** *Slc32a1*, **f** *Slc17a6*, **g** *Crh*, **h** *Prkcd*, **i** *Glp1r*, **j** *Th,* and **k** *Esr1*. Scale bar, 20 µm. **l, m** Percentage of *Mc3r* neurons expressing *Slc32a1*, *Slc17a6*, *Crh*, *Prkcd*, *Glp1r*, *Th*, and *Esr1*. *Mc3r* neurons exhibit the highest coexpression with *Slc32a1*. aco, anterior commissure; al, anterolateral; am, anteromedial; BST, bed nuclei of the stria terminalis; Crh, corticotropin-releasing hormone; Esr1, estrogen receptor 1; fx, fornix; fu, fusiform; Glp1r, glucagon-like peptide-1 receptor; ju, juxtacapsular; mg, magnocellular; ov, oval; pr, principal; Prkcd, protein kinase C delta; rh, rhomboid; Slc17a6, solute carrier family 17 member 6 (vesicular glutamate transporter 2 gene); Slc32a1, solute carrier family 32 member 1 (vesicular GABA Transporter gene); Th, tyrosine hydroxylase.

In addition to mapping the distribution of MC3R neurons throughout the BST, we also aimed to determine if MC3R neuronal labeling in the BSTd is significantly different between males and females. No differences in the number of labeled neurons were found between male and female MC3R-Cre::tdTomato mice in the BSTd (Figure 1d).

To gain further understanding of the overlap of MC3R neurons in the BSTd with other neural substrates coordinating feeding, stress and anxiety-like behaviors, multiplex fluorescent *in situ* hybridization was utilized (Figure 1e-m). No sex differences in coexpression were observed with any of the substrates examined and thus male and female data were combined. *Mc3r* neurons exhibited the highest coexpression with *Slc32a1,* the gene that codes for vesicular inhibitory amino acid transporter, VGAT, (86.9 ± 6.0%; Figure 1e,l,m). *Mc3r* neurons also highly coexpressed substrates known to modulate feeding and anxiety-like behaviors, including *Prkcd* (72.8 ± 5.7%; Figure 1h,l,m), and *Glp1r* (71.0 ± 8.3%; Figure 1i,l,m). *Mc3r* neurons had moderate to high overlapping expression with *Esr1* (65.3 ± 6.4%; Figure 1k,l,m), *Crh* (58.7 ± 8.0%; Figure 1g,l,m) and *Th* (50.6 ± 12.1%; Figure 1j,l,m). Low coexpression was detected with *Slc17a6*, the gene that codes for vesicular glutamate transporter 2, VGLUT2, (9.6 ± 1.6%; Figure 1f,l,m). Therefore, BST^MC3R^ neurons are mainly GABAergic and are comprised of subpopulations of peptidergic neurons thought to drive feeding, stress and anxiety-like behaviors.

### Activation of BST^MC3R^ Neurons Influences Feeding and Responses to Stress but not Anxiety-Like Behavior

To determine if activating BST^MC3R^ neurons impacts feeding, stress and anxiety-like behaviors, a chemogenetic approach was used. An adeno-associated virus (AAV) expressing the Cre-dependent excitatory DREADD (designer receptors exclusively activated by designer drugs) activator, hM3Dq, was injected into the BSTd of male and female MC3R-Cre mice (Figure 2a). Histological validation revealed targeted hM3Dq-mCherry labeling in the BSTd like what was observed in MC3R-Cre::tdTomato mice (Figure 2b). In addition to hM3Dq injections, additional control groups of male and female mice were injected with AAV5-hSyn-DIO-mCherry to ensure CNO did not have off-target effects. No differences were observed between mCherry control animals that received saline or CNO in any behavior recorded (Figure S1).

**Figure 2.**
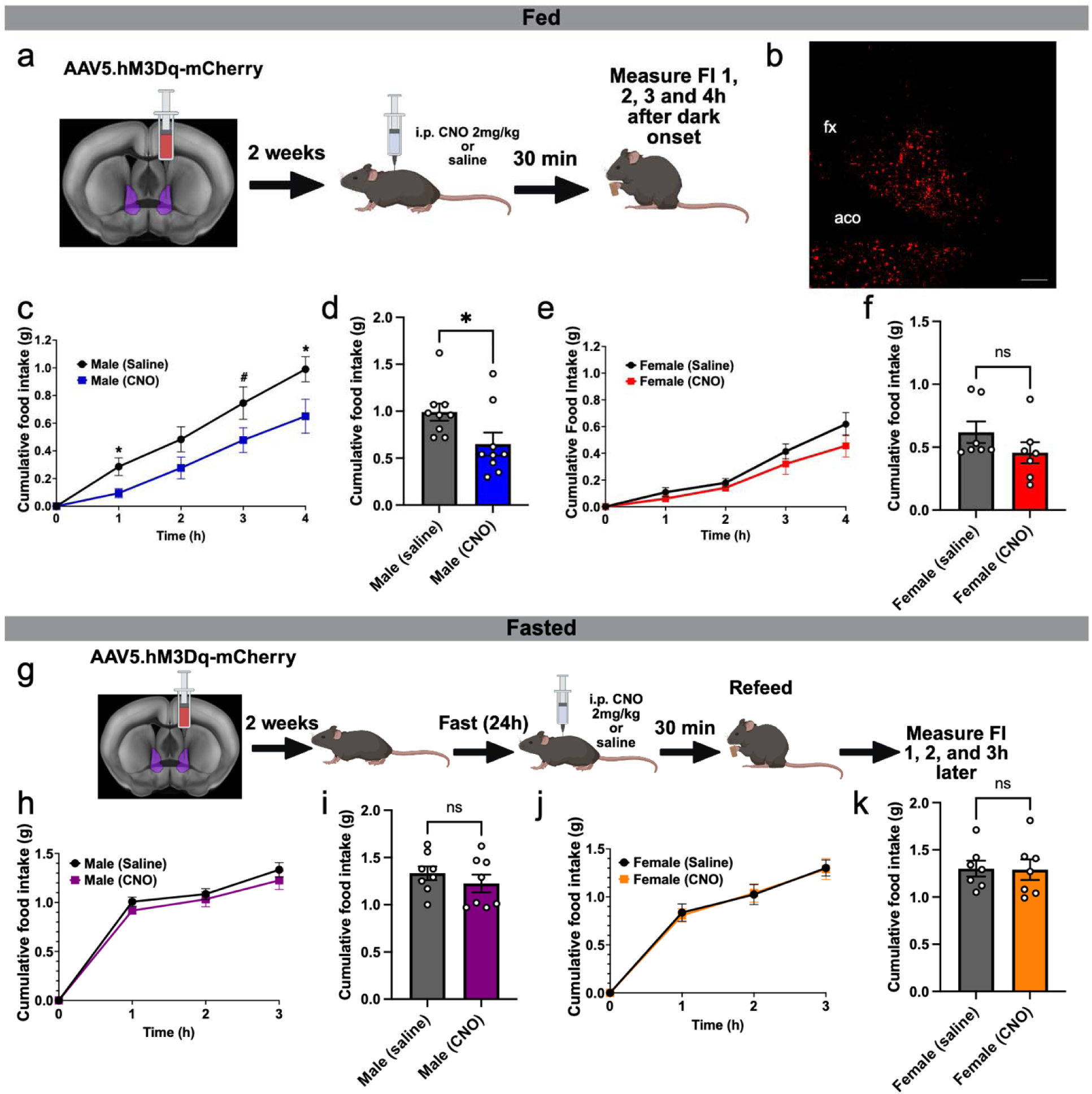
BST^MC3R^ neurons decrease feeding in the fed state in males. **a** Schematic of experimental paradigm. AAV5.hM3Dq-mCherry was injected into the BSTd of male and female MC3R-Cre mice. Animals were allowed to recover for at least two weeks prior to any experiments being conducted. On the day of the experiment, mice received an injection of either CNO (2mg/kg) or saline 30 minutes prior to access to food that began at the start of the dark phase. Food intake was measured for 4 hours. **b** Low magnification confocal image depicting representative injection site of AAV5.hM3Dq- mCherry. Scale bar, 200 µm. **c-f** Chemogenetically exciting BST^MC3R^ neurons in MC3R-Cre mice in a fed state blunts cumulative 4 hour feeding in **c, d** males but not **e, f** females (c, e: n=7-10/group, 2-way ANOVA, Sidaks multiple comparisons test; d, f: n=7-10/group, unpaired t-test). **g** Schematic of experimental paradigm. AAV5.hM3Dq-mCherry was injected into the BSTd of male and female MC3R-Cre mice. Animals were allowed to recover for at least two weeks prior to any experiments being conducted. Mice were fasted for 24 hours. On the day of the experiment, mice received an injection of either CNO (2mg/kg) or saline 30 minutes prior to access to food. Food intake was measured for 3 hours. **h-k** Chemogenetically exciting BST^MC3R^ neurons in MC3R-Cre mice has no significant effect on refeeding after a fast in **h, i** males or **j, k** females (h, j: n=7-8/group, 2-way ANOVA, Sidaks multiple comparisons test; i, k: n=7-8/group, unpaired t-test). Aco, anterior commissure; fx, fornix. * p < 0.05, # p < 0.09.

To define the impact of BST^MC3R^ neuron chemogenetic activation on feeding during the fed state, CNO or saline was injected 30 min prior to access to food that began at the start of the dark phase in the home cage of the mouse. Food intake was subsequently recorded for 4 hours. Chemogenetically stimulating BST^MC3R^ neurons significantly decreased food intake in males in a fed state (Figure 2c,d). In contrast, stimulating BST^MC3R^ neurons in females in a fed state did not have a significant effect on food intake (Figure 2e,f).

In addition to elucidating if chemogenetic activation of BST^MC3R^ neurons influences feeding in the fed state, we also examined if chemogenetic activation of BST^MC3R^ neurons could alter feeding in mice in a fasted state. Male and female mice were fasted for 24 hours, given an injection of CNO or saline 30 min prior to access to food, and food intake was measured for the next 3 hours (Figure 2g). Unlike what was observed in the fed state, chemogenetically activating MC3R neurons in the fasted state had no effect on food intake in male or female mice (Figure 2h-k). Thus, MC3R activation seems to be particularly important for decreasing feeding in males in a fed state.

We next tested whether chemogenetic activation of BST^MC3R^ neurons would affect anxiety-like behaviors. In the open field test (OFT) and elevated plus maze (EPM), chemogenetically activating BST^MC3R^ neurons had no measurable effect on anxiety-like behaviors (Figure 3a-h). In addition to identifying if activation of BST^MC3R^ neurons would affect behaviors in general anxiety-like behavior tests, we also wanted to determine if BST^MC3R^ neurons modulated anxiety-like behaviors in a stress-induced anorexia paradigm. To do this, we performed the novelty-suppressed feeding test (NSFT) in which male and female mice were fasted for 24 hours, placed into a novel environment, and a single pellet of normal chow was placed in the center of the open field (Figure 3i). Chemogenetically activating BST^MC3R^ neurons had no effect on cumulative food intake (Figure 3j,k) or the latency to feed (Figure 3l,m) in male or female mice. Therefore, chemogenetic activation of BST^MC3R^ neurons at baseline does not seem to influence anxiety-like behaviors.

**Figure 3.**
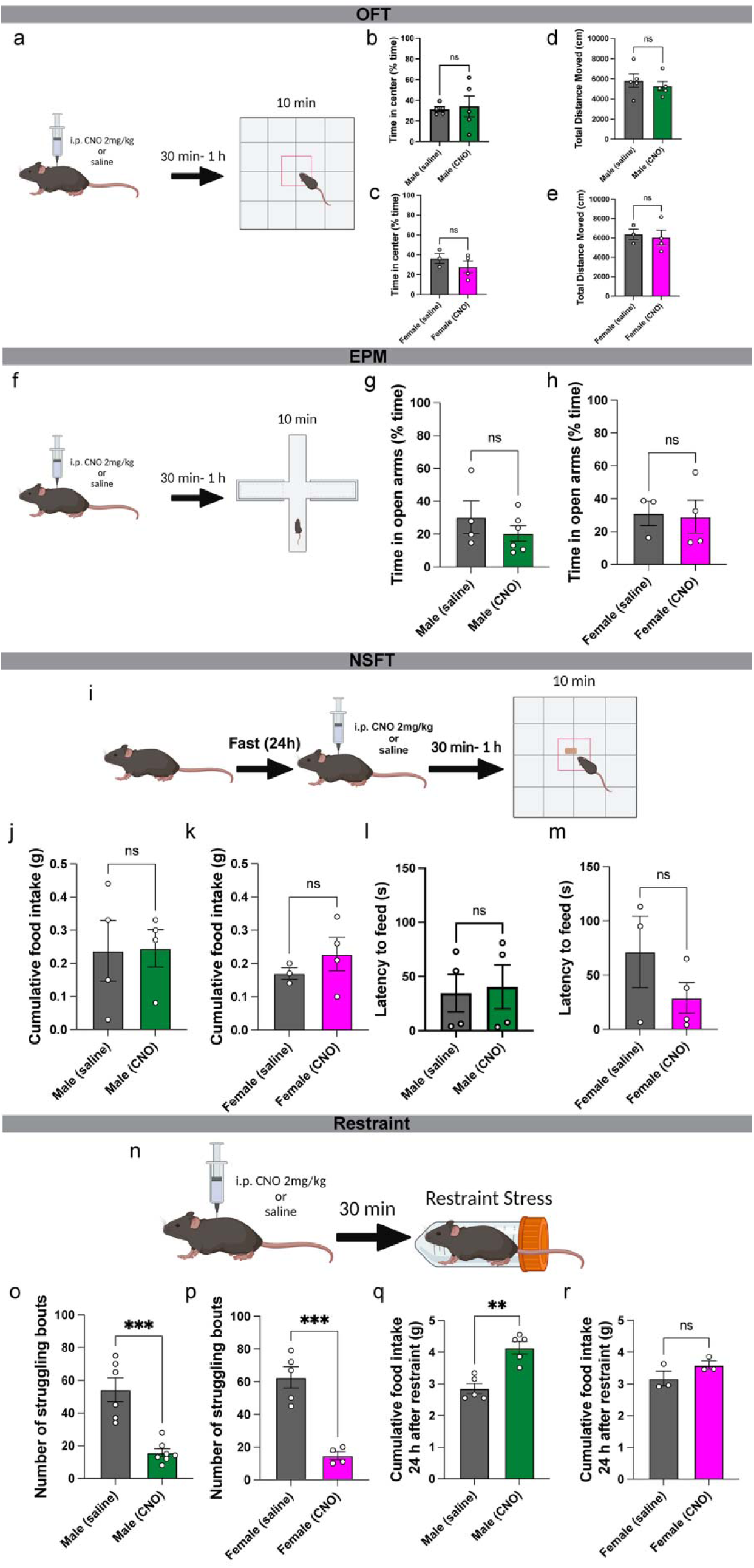
BST^MC3R^ neurons decrease responses to stress but have no effect on anxiety-like behavior. **a** Schematic of open field test (OFT) experimental paradigm. On the day of the experiment, mice received an injection of either CNO (2mg/kg) or saline 30 minutes prior to being placed in the apparatus. The behavior of male and female MC3R-Cre mice in the open field was recorded for 10 minutes. **b, c** Chemogenetically exciting BST^MC3R^ neurons in MC3R-Cre mice has no significant effect on the percentage of time spent in the center of the OFT in **b** males or **c** females (n=3-5/group, unpaired t-test). **d,e** Chemogenetically exciting BST^MC3R^ neurons in MC3R-Cre mice has no significant effect on total distance moved during the OFT in **d** males or **e** females (n=3-5/group, unpaired t-test). **f** Schematic of elevated plus maze (EPM) experimental paradigm. On the day of the experiment, mice received an injection of either CNO (2mg/kg) or saline 30 minutes prior to being placed in the apparatus. The behavior of male and female MC3R-Cre mice in the EPM was recorded for 10 minutes. **g, h** Chemogenetically exciting BST^MC3R^ neurons in MC3R-Cre mice has no significant effect on the percentage of time spent in the open arms of the EPM in **g** males or **h** females (n=3-6/group, unpaired t-test). **i** Schematic of the novelty-suppressed feeding test experimental paradigm. Mice were fasted for 24 h. On the day of the experiment, mice received an injection of either CNO (2mg/kg) or saline 30 minutes prior to being placed in the apparatus. The behavior of male and female MC3R-Cre mice in the apparatus was recorded for 10 minutes. **j-m** Chemogenetically exciting BST^MC3R^ neurons in MC3R-Cre mice has no significant effect on cumulative food intake or latency to feed in **j, l** males or **k, m** females during the NSFT (n=3-4/group, unpaired t-test). **n** Schematic of restraint stress experimental paradigm. On the day of the experiment, mice received an injection of either CNO (2mg/kg) or saline 30 minutes prior to being placed in a conical tube. The struggling behavior of male and female MC3R-Cre mice was recorded for 30 minutes. **o, p** Chemogenetically exciting BST^MC3R^ neurons in MC3R-Cre mice significantly decreases the number of struggling bouts in **o** males and **p** females during restraint stress (n=4-7/group, unpaired t-test). **q, r** Chemogenetically exciting BST^MC3R^ neurons during restraint significantly increases cumulative food intake in the 24 hours after restraint in **q** males but not **r** females (n=3-5/group, unpaired t-test). ** p < 0.01, *** p < 0.001.

However, BST^MC3R^ neurons could play a role in responses to other types of stressors. To examine this, male and female mice were given an injection of CNO 30 minutes prior to being restrained. The number of struggling bouts, a previously validated measure of stress coping^35,45^, that occurred during a 30-minute period was recorded (Figure 3n). In both male and female mice, chemogenetically activating BST^MC3R^ neurons significantly decreased the number of struggling bouts (Figure 3o,p). In addition to measuring struggling bouts, we also measured the amount of food mice ate cumulatively in their home cages in the 24 hours after restraint stress to investigate if changing the response to stress would change the typical decrease in food intake after restraint stress^46^. Indeed, chemogenetically stimulating BST^MC3R^ neurons prior to restraint stress significantly increased food intake after restraint in males (Figure 3q) but not females (Figure 3r). Altogether, chemogenetic activation of BST^MC3R^ neurons decreases the response to restraint stress in both sexes but protects against stress-induced anorexia only in males.

### BST^MC3R^ Neurons are Activated During Feeding and Responses to Stress

Because chemogenetically activating BST^MC3R^ neurons impacted feeding behavior and responses to stress, we next wanted to explore if BST^MC3R^ neuronal activity is time-locked to these behaviors using *in vivo* fiber photometry in freely behaving mice. An AAV expressing the Cre-dependent genetically encoded calcium indicator, GCaMP7f, was injected into the BSTd of male and female MC3R-Cre mice, and a fiberoptic cannula was implanted immediately following injection (Figure 4a). Histological validation revealed targeted GCaMP labeling in the BSTd like what was observed in MC3R-Cre::tdTomato mice (Figure 4b).

**Figure 4.**
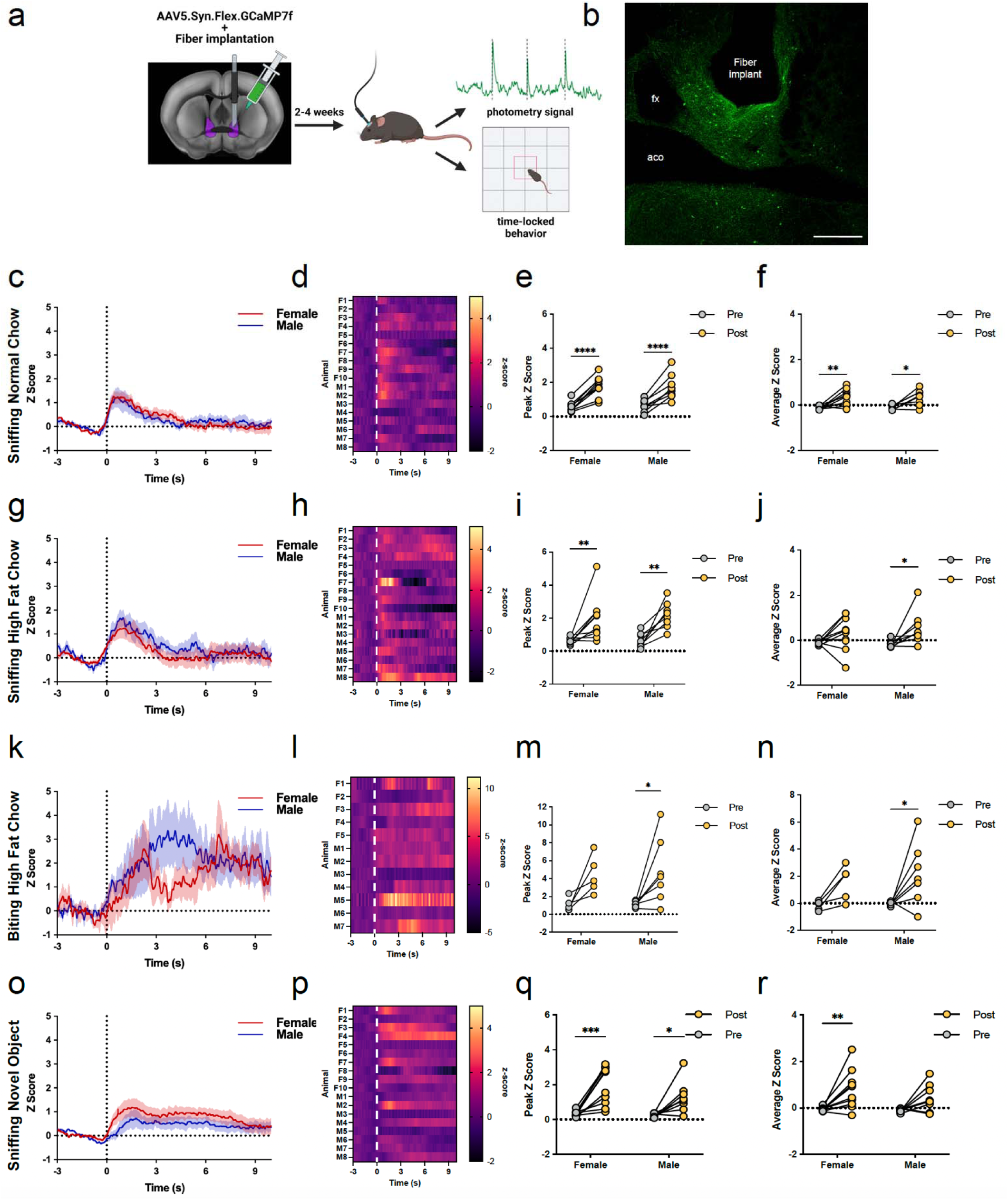
BST^MC3R^ neurons are activated during feeding behaviors in a fed state. **a** Schematic of experimental paradigm. AAV5.Syn.Flex.GCaMP7f was injected into the BSTd of male and female MC3R-Cre mice and a fiberoptic cannula was implanted immediately after injection. Animals were allowed to recover for at least two weeks prior to any experiments being conducted. On the day of the experiment, photometry signal and animal behavior was concurrently recorded to time-lock GCaMP signal with specific behaviors. **b** Low magnification confocal image depicting representative injection site of AAV5.Syn.Flex.GCaMP7f and fiberoptic cannula placement. Scale bar, 500 µm. **c** Average time-locked GCaMP signal while sniffing a normal chow pellet in males (blue) and females (red). **d** Individual calcium transient traces of the average z-score for each animal. **e, f** Both the **e** peak and **f** average z-score is significantly greater than baseline in both males and females (n=8-10/group, 2-way ANOVA, Sidaks multiple comparisons test). **g** Average time-locked GCaMP signal while sniffing a high fat chow pellet in males (blue) and females (red). **h** Individual calcium transient traces of the average z-score for each animal **i** The peak z-score is significantly greater than baseline in both males and females (n=8-10/group, 2-way ANOVA, Sidaks multiple comparisons test). **j** The average z-score is significantly greater in males only (n=8-10/group, 2-way ANOVA, Sidaks multiple comparisons test). **k** Average time-locked GCaMP signal while biting a high fat chow pellet in males (blue) and females (red). **l** Individual calcium transient traces of the average z-score for each animal. **m, n** Both the **m** peak and **n** average z-score is significantly greater than baseline in males only (n=8-10/group, 2-way ANOVA, Sidaks multiple comparisons test). **o** Average time-locked GCaMP signal while sniffing a novel object (battery) in males (blue) and females (red). **p** Individual calcium transient traces of the average z-score for each animal. **q** The peak z-score is significantly greater than baseline in both males and females (n=8-10/group, 2-way ANOVA, Sidaks multiple comparisons test). **r** The average z-score is significantly greater in females only (n=8-10/group, 2-way ANOVA, Sidaks multiple comparisons test). Aco, anterior commissure; fx, fornix. * p < 0.05, ** p < 0.01. *** p < 0.001, **** p < 0.0001.

In a fed state, BST^MC3R^ neuronal activity was significantly increased while both male and female mice approached and sniffed a normal chow (Figure 4c-f) and high fat chow (Figure 4g-j) pellet. BST^MC3R^ neuronal activity was also increased when mice interacted with a novel object that was not food (Figure 4o-r), but the peak response (Figure 4q) was markedly less than when sniffing high fat chow (Figure 4i). The largest increase in BST^MC3R^ neuronal activity in a fed state occurred when mice were biting a high fat chow pellet (Figure 4k-n). Interestingly, this increase in activity was only significantly elevated from baseline in males (Figure 4k-n). Mice did not bite a normal chow pellet while in a fed state and thus BST^MC3R^ neuronal activity was not aligned to this event. Altogether, BST^MC3R^ neurons are activated when mice are interacting with both food and non-food objects with the most increased activation occurring while males specifically eat a palatable diet. Therefore, BST^MC3R^ neurons may play a particularly important role in hedonic feeding in a sexually dimorphic manner.

In comparison to the fed state, BST^MC3R^ neurons were not as responsive to food in a fasted state (Figure 5). Unlike in the fed state, the average z-score did not significantly differ from baseline in any of the behaviors analyzed in male or female mice (Figure 5d,h,l,p). The peak response of BST^MC3R^ neuronal activity was significantly increased when sniffing normal chow in females (Figure 5c), as well as while biting normal and high fat chow in males (Figure 5k,o).

**Figure 5.**
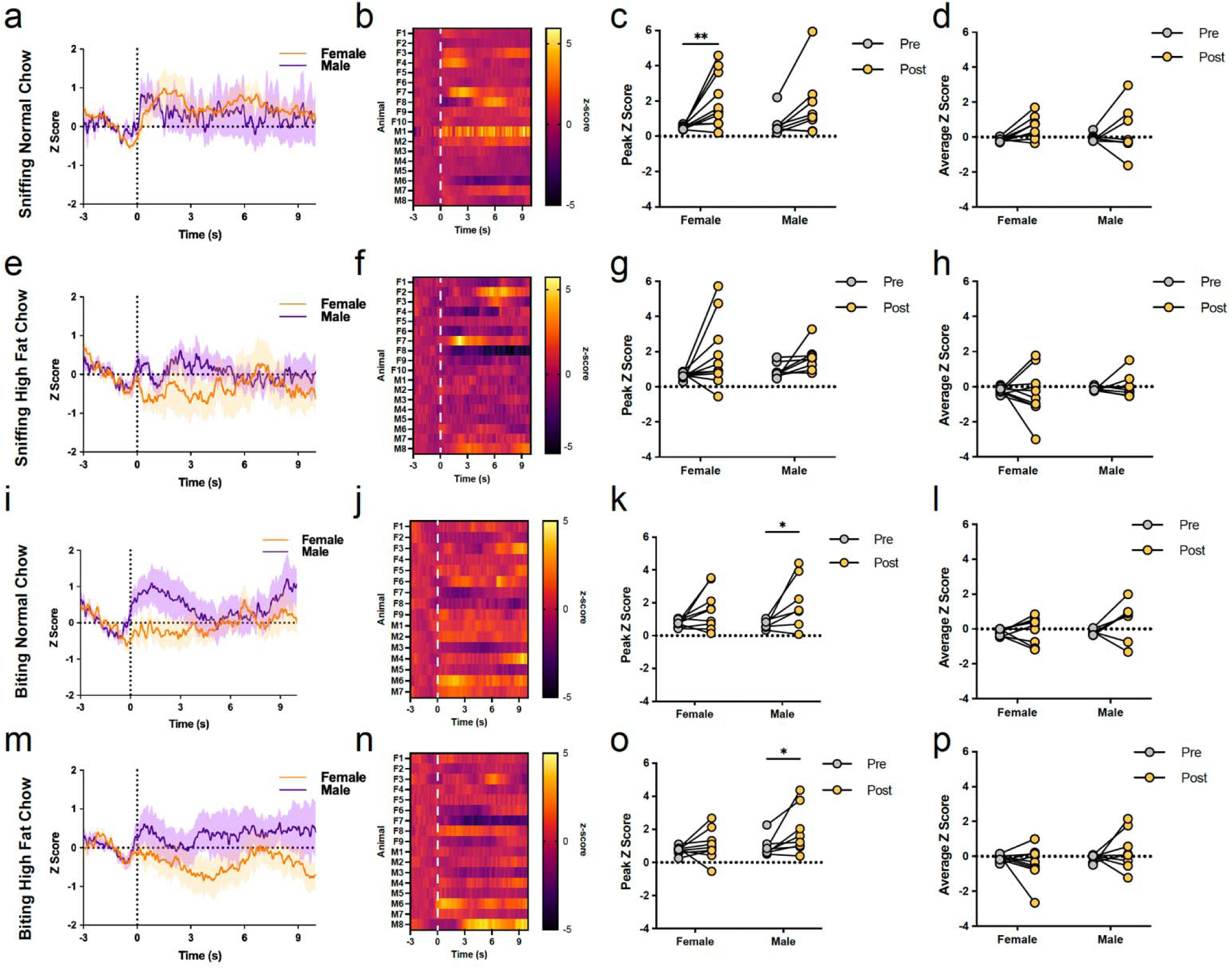
BST^MC3R^ neurons are minimally activated during feeding behaviors in a fasted state. **a** Average time-locked GCaMP signal while sniffing a normal chow pellet in males (purple) and females (orange). **b** Individual calcium transient traces of the average z-score for each animal. **c** The peak z-score is significantly greater than baseline in females only (n=8-10/group, 2-way ANOVA, Sidaks multiple comparisons test). **d** The average z-score is not significantly greater than baseline in males or females (n=8-10/group, 2-way ANOVA, Sidaks multiple comparisons test). **e** Average time-locked GCaMP signal while sniffing a high fat chow pellet in males (purple) and females (orange). **f** Individual calcium transient traces of the average z-score for each animal. **g, h** Neither the **g** peak nor the **h** average z-score is significantly different from baseline in males or females (n=8-10/group, 2-way ANOVA, Sidaks multiple comparisons test). **i** Average time-locked GCaMP signal while biting a normal chow pellet in males (purple) and females (orange). **j** Individual calcium transient traces of the average z-score for each animal. **k** The peak z-score is significantly greater than baseline in males only (n=8-10/group, 2-way ANOVA, Sidaks multiple comparisons test). **l** The average z-score is not significantly greater than baseline in males or females (n=8-10/group, 2-way ANOVA, Sidaks multiple comparisons test). **m** Average time-locked GCaMP signal while biting a high fat chow pellet in males (purple) and females (orange). **n** Individual calcium transient traces of the average z-score for each animal. **o** The peak z-score is significantly greater than baseline in males only (n=8-10/group, 2-way ANOVA, Sidaks multiple comparisons test). **p** The average z-score is not significantly greater than baseline in males or females (n=8-10/group, 2-way ANOVA, Sidaks multiple comparisons test). * p < 0.05, ** p < 0.01.

In addition to responding to food, BST^MC3R^ neurons were also activated during defensive behaviors, exploratory behaviors, and responses to stress (Figure 6). When faced with visual stimuli that mimics a rapidly approaching aerial predator (looming), BST^MC3R^ neuronal activity significantly increased in male and female mice (Figure 6a-d). BST^MC3R^ neuronal activity also significantly increased when mice rear on the side of the cage to explore their surroundings (Figure 6e-h). Finally, we exposed the mice to three different speeds of shaking to determine if BST^MC3R^ neuronal activity is increased when exposed to a stressor that escalates in intensity (Figure 6i-t). Neither male nor female mice experienced a significant increase in BST^MC3R^ neuronal activity during slow shaker stress (Figure 6i-l). However, BST^MC3R^ neuronal activity was significantly increased during moderate (Figure 6m-p) and high (Figure 6q-t) shaker stress. Taken together, BST^MC3R^ neurons are significantly activated, and may play a role, in defensive, exploratory and stressful behaviors.

**Figure 6.**
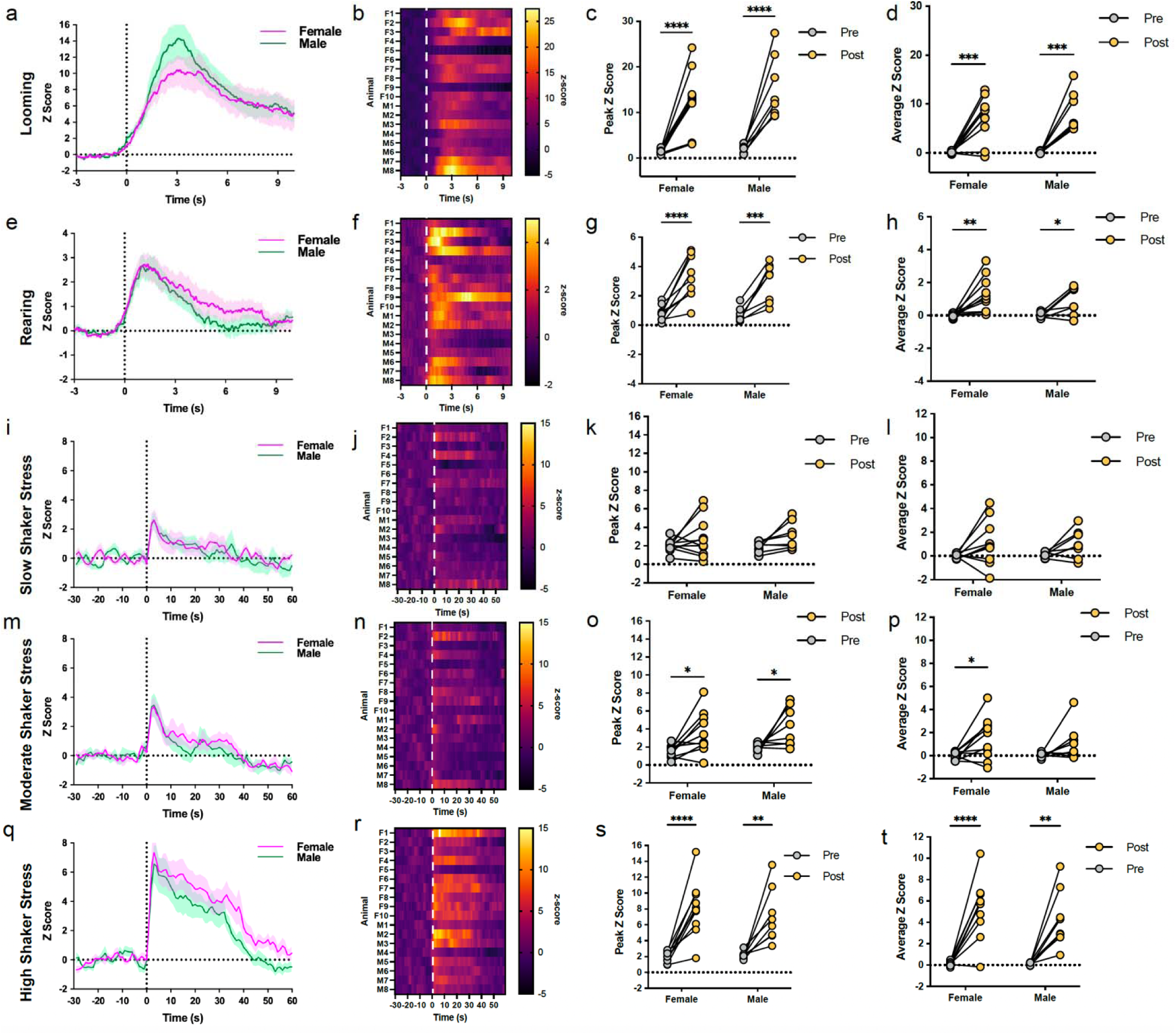
BST^MC3R^ neurons are activated during defensive behaviors, exploratory behaviors and in response to stressors. **a** Average time-locked GCaMP signal during looming behavior in males (green) and females (pink). **b** Individual calcium transient traces of the average z-score for each animal. **c, d** The **c** peak and **d** average z-score is significantly greater than baseline in males and females (n=8-10/group, 2-way ANOVA, Sidaks multiple comparisons test). **e** Average time-locked GCaMP signal during rearing behavior in males (green) and females (pink). **f** Individual calcium transient traces of the average z-score for each animal. **g, h** The **g** peak and **h** average z-score is significantly greater than baseline in males and females (n=8-10/group, 2-way ANOVA, Sidaks multiple comparisons test). **i** Average time-locked GCaMP signal during slow shaker stress in males (green) and females (pink). **j** Individual calcium transient traces of the average z-score for each animal. **k, l** Neither the **k** peak nor the **l** average z-score is significantly greater than baseline in males and females (n=8-10/group, 2-way ANOVA, Sidaks multiple comparisons test). **m** Average time-locked GCaMP signal during moderate shaker stress in males (green) and females (pink). **n** Individual calcium transient traces of the average z-score for each animal. **o** The peak z-score is significantly greater than baseline in males and females (n=8-10/group, 2-way ANOVA, Sidaks multiple comparisons test). **p** The average z-score is significantly greater than baseline in females only (n=8-10/group, 2-way ANOVA, Sidaks multiple comparisons test). **q** Average time-locked GCaMP signal during fast shaker stress in males (green) and females (pink). **r** Individual calcium transient traces of the average z-score for each animal. **s, t** The **s** peak and **t** average z-score is significantly greater than baseline in males and females (n=8-10/group, 2-way ANOVA, Sidaks multiple comparisons test). * p < 0.05, ** p < 0.01. *** p < 0.001, **** p < 0.0001.

### BST^MC3R^ Neurons Send Projections to Brain Regions That Drive Feeding and Responses to Stress

Since BST^MC3R^ neurons are activated during, and can stimulate, feeding and responses to stress, we wanted to ascertain the downstream targets of BST^MC3R^ neurons that could play a role in driving these behaviors. We injected MC3R-Cre mice with an AAV expressing the Cre-dependent anterograde tracer, Synaptophysin-mRuby, into the BSTd (Figure 7a). BST^MC3R^ neurons form dense terminal fields in multiple brain regions that influence several different classes of motivated behavior, including ingestive, affective, reproductive, defensive and arousal behaviors, among others (Figure 7b-n). In the striatum, the nucleus accumbens core exhibited profuse axonal labeling (Figure bB). Several regions of the hypothalamus contained a moderate to high density of labeled fibers, including the MEPO (Figure 7c), LPO (Figure 7c), MPO (Figure 7c,d), AVPV (Figure 7d), MPN (Figure 7f), PVpo (Figure 7f), AHN (Figure 7g), PVH (Figure 7g), LHA (Figure 7h,i), DMH (Figure 7i), and VMHvl (Figure 7j). Dense terminal fields were also observed in the amygdala with the highest density of inputs localized to the MEA (Figure 7k). Light to moderate densities of labeled axons were present in the medial and lateral septum (Figure 7e). In the midbrain, the PAG (Figure 7m) exhibited moderately dense terminal labeling, whereas the VTA (Figure 7l) and PBmm (Figure 7n) contained sparser (but notable) labeled inputs. In addition to mapping the downstream targets of BST^MC3R^ neurons, we also wanted to determine if BST^MC3R^ terminal densities were sexually dimorphic in regions known to play a role in feeding and responses to stress, which could explain the sexually dimorphic behaviors observed in chemogenetic and fiber photometry experiments. Surprisingly, we did not observe marked sex differences in these terminal fields. To test this quantitatively, we measured the density of BST^MC3R^ axonal labeling in the ACB, LHA, PAG, and VTA in males and females (Figure 7o). There were no sex differences in the density of axonal labeling in any of the areas examined (Figure 7o). Therefore, while BST^MC3R^ neurons send dense projections to multiple brain nuclei that drive feeding behaviors and responses to stress, differences between males and females were not observed that would explain the sexually dimorphic behaviors discovered in chemogenetic and fiber photometry experiments.

**Figure 7.**
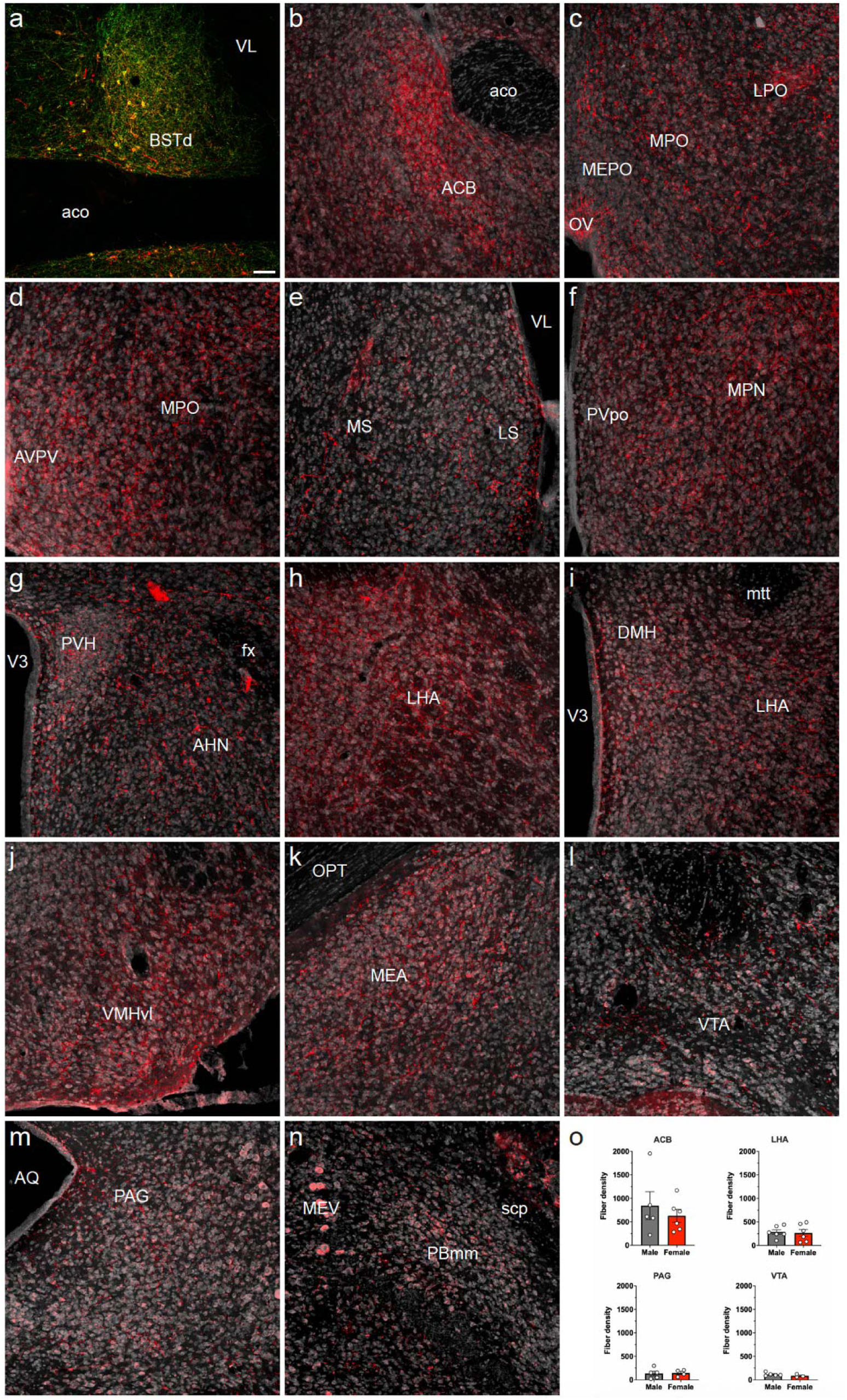
BST^MC3R^ neurons send dense projections to several brain regions that drive feeding behaviors, defensive behaviors and responses to stress. **a** Low magnification confocal image depicting representative injection site of AAVDJ-hSyn-FLEx-mGFP-2A-Synaptophysin-mRuby in the BST of an MC3R-cre mouse. Scale bar, 100 µm. **b-n** BST^MC3R^ neuronal projections are present in the **b** ACB, **c** LPO, **c** MEPO, **c, d** MPO, **e** LS, **e** MS, **f** MPN, **f** PVpo, **g** AHN, **g** PVH, **h, i** LHA, **i** DMH, **j** VMHvl, **k** MEA, **l** VTA, **m** PAG, and **n** PBmm. **o** There were no sex differences in terminal density in any of the areas measured (n=3-6/group, unpaired t-test). ACB, nucleus accumbens; aco, anterior commissure; AHN, anterior hypothalamic nucleus; AVPV, anteroventral periventricular nucleus; AQ, cerebral aqueduct; BSTd, dorsal bed nuclei of the stria terminalis; DMH, dorsomedial hypothalamic nucleus; fx, fornix; LHA, lateral hypothalamic area; LPO, lateral preoptic area; LS, lateral septal nucleus; MEA, medial amygdalar nucleus; MEPO, median preoptic nucleus; MEV, midbrain trigeminal nucleus; ML, medial septal nucleus; MPN, medial preoptic nucleus; MPO, medial preoptic area; mtt, mammillothalamic tract; OPT, optic tract; OV, vascular organ of the lamina terminalis; PBmm, parabrachial nucleus, medial division, medial medial part; PAG, periaqueductal gray; PVH, paraventricular hypothalamic nucleus; PVpo, periventricular hypothalamic nucleus, preoptic part; scp, superior cerebellar peduncles; V3, third ventricle; VL, lateral ventricle; VMHvl, ventromedial hypothalamic nucleus, ventrolateral part; VTA, ventral tegmental area.

### Brain Regions That Modulate Feeding and Responses to Stress Provide Monosynaptic Inputs to BST^MC3R^ Neurons

Although BST^MC3R^ projections do not appear to be sexually dimorphic, the sex differences in behavioral responses we observed may be due to sexually dimorphic afferents. To address this question, we employed a rabies-mediated tract-tracing approach, in combination with tissue clearing and whole brain lightsheet imaging to create a quantitative brain-wide map of neurons providing monosynaptic inputs to BST^MC3R^ neurons (Figure 8a). The distribution of labeled neurons suggests that BST^MC3R^ neurons receive dense monosynaptic inputs from brain regions known to participate in relaying cognitive, emotional, autonomic and neuroendocrine information (Figure 8b-h). Multiple nuclei in the hypothalamus exhibited high densities of afferents to BST^MC3R^ neurons, as over half of the top 50 densest regions originate from this area (Figure 8h). A significant density of neurons in the isocortex, olfactory areas, striatum, pallidum, thalamus and medulla also send projections to BST^MC3R^ neurons (Figure 8h). There was no region where one sex received afferents that the other did not. However, several differences in the density of neurons projecting to BST^MC3R^ neurons were found between males and females (Figure 8i-n). BST^MC3R^ neurons in males received an increased density of afferents from the parasubthalamic nucleus, parataenial nucleus, paraventricular nucleus of the thalamus, nucleus accumbens, nucleus of reuniens and medial group of the dorsal thalamus compared to females (Figure 8n). In contrast, females received an increased density of afferents from the preoptic part of the periventricular hypothalamic nucleus, medial septal nucleus and periventricular zone compared to males (Figure 8n). Therefore, the density of several afferents to BST^MC3R^ neurons is sexually dimorphic and may contribute to expression of sexually dimorphic feeding behaviors and responses to stress.

**Figure 8.**
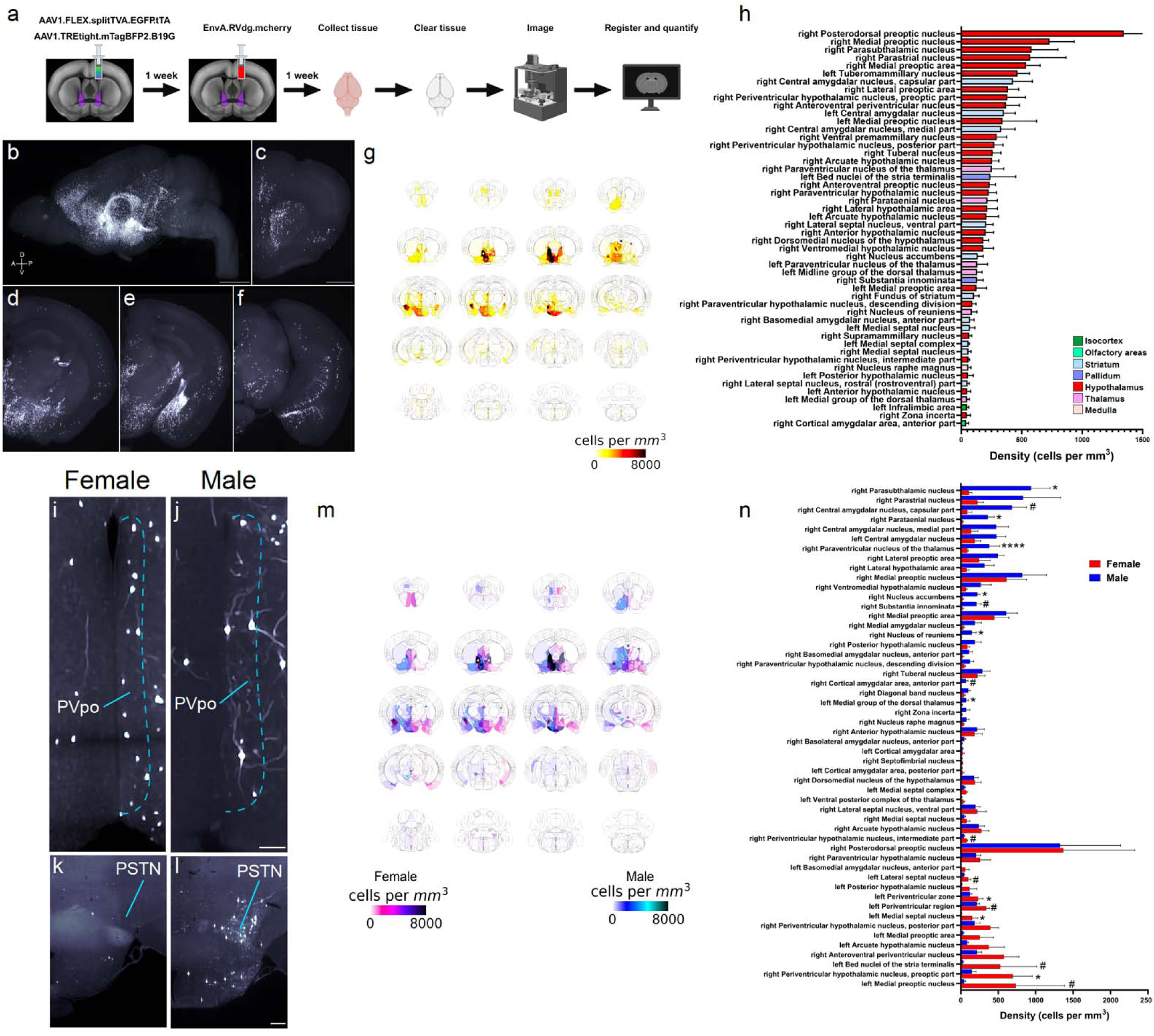
Monosynaptic inputs to BST^MC3R^ neurons originate from brain regions relaying cognitive, emotional, neuroendocrine and autonomic information and are sexually dimorphic. **a** Schematic of whole brain analysis pipeline. AAV1.FLEX.splitTVA.EGFPtTA and AAV1.TREtight.m.TagBFP2.B19G were injected into the right BSTd of male and female MC3R-Cre mice. One week later, EnvA-RVdg.mcherry was injected into the same area. Tissue was collected one week later, cleared, imaged and registered to the Allen Common Coordinate Reference Framework. **b** Three-dimensional rendering of a unilateral monosynaptic rabies injection into the BST of an MC3R-cre mouse. Scale bar, 1mm. **c-f** Coronal view of monosynaptic inputs to BST^MC3R^ neurons in the same brain at the level of the **c** prelimbic and insular cortex, **d** paraventricular thalamus and hypothalamus, **e** lateral hypothalamic area and central amygdala, and **f** hippocampus and periaqueductal gray. Scale bar, 500 µm. **g** Average cell density of all animals (n=7) quantified by registering cell locations to the Allen Common Coordinate Reference Framework. **h** Top 50 areas of total average cell density in all animals (n=7) quantified by registering cell locations to the Allen Common Coordinate Framework. Over half of the top 50 areas that send inputs to BST^MC3R^ neurons originate in the hypothalamus. **i, j** Representative images of rabies labeling in the PVpo of a **i** female and **j** male MC3R-cre mouse. Females have significantly more PVpo neurons sending inputs to BST^MC3R^ neurons than males (n=3-4/group, unpaired t-test). Scale bar, 100 µm. **k, l** Representative images of rabies labeling in the PSTN of a **k** female and **l** male MC3R-cre mouse. Males have significantly more PSTN neurons sending inputs to BST^MC3R^ neurons than females (n=3-4/group, unpaired t-test). Scale bar, 100 µm. **m, n** Comparison of average cell density between females and males quantified by registering cell locations to the Allen Common Coordinate Reference Framework (n=3-4/group, unpaired t-test). PSTN, parasubthalamic nucleus; PVpo, periventricular hypothalamic nucleus, preoptic part. # p < 0.06, * p < 0.05, **** p < 0.0001.

## DISCUSSION

Previous work revealed MC3R neurons bidirectionally regulate feeding and anxiety-like behaviors, and global deletion of MC3R produces multiple forms of sexually dimorphic anxiety-related hypophagia^32^. Our findings support the emerging notion that the BST^MC3R^ circuitry lies at the intersection of feeding, stress and anxiety-like behaviors^33,47–49^ and identify BST^MC3R^ neurons as a key population of neurons serving this function. An examination of MC3R neuronal subtypes in the BSTd found that they are primarily GABAergic and comprise molecularly heterogeneous subpopulations that may impact feeding, stress and anxiety-like behaviors differently in males and females. BST^MC3R^ neurons are activated during feeding behaviors, and chemogenetic stimulation of BST^MC3R^ neurons induces decreases in feeding, particularly in the fed state in males. BST^MC3R^ neurons are also significantly activated and can influence responses to stress and defensive behaviors in both males and females. BST^MC3R^ neurons send dense projections to multiple brain nuclei known to coordinate feeding behaviors and responses to stress, but these projections appear to be similar in males and females suggesting that the sexually dimorphic behaviors discovered in chemogenetic and fiber photometry experiments are not likely to be due to sex differences in the pattern of BST^MC3R^ neuronal projections. In contrast, afferents to BST^MC3R^ neurons from multiple regions are sexually dimorphic. In females, high densities of afferent neurons are found in regions known to play a role in social and maternal behaviors, including the medial preoptic area, periventricular preoptic region, medial and lateral septum and periventricular zone. Conversely, a higher density of afferents from regions involved in feeding, stress, anxiety and autonomic regulation, including the parasubthalamic nucleus, central amygdala, paraventricular hypothalamus, nucleus accumbens, substantia inominata and nucleus of reuniens are observed in males. This increased afferent input in males suggests there may be fundamental differences in how metabolic information is integrated with cognitive, emotional and autonomic information in males and females that could be driving sexually dimorphic feeding behaviors and responses to stress. Altogether, these data reveal that afferents, but not projections, of BST^MC3R^ neurons are sexually dimorphic and that these cells may be involved in sexually dimorphic aspects of feeding, responses to stress and defensive behaviors.

Several neuropeptides and transmitters within the BST have been implicated in the control of feeding, energy homeostasis, anxiety and stress ^5,12–17,23^. Here, we found that MC3R neurons in the BSTd are primarily GABAergic, but also express *Crh*, *Glp1r*, *Esr1*, *Th* and *Prkcd* with *Prkcd* being the most highly coexpressed transcript. However, it is important to note that *Prkcd* is mainly expressed in BSTov, where there are significantly more cells expressing *Prkcd* than *Mc3r, and* only 20% of these coexpress Mc3r (unpublished data). A substantial number of *Mc3r* neurons also coexpress *Crh* and *Glp1r*. BST^Crh^ neurons are known to modulate unconditioned defensive responses^50^, responses to acute stress^51^, negative affective behaviors^35–37,52^ and energy homeostasis^53^, whereas BST^GLP1R^ neurons are known to decrease feeding^46^. These effects are comparable to responses we observed in chemogenetic and fiber photometry experiments with BST^MC3R^ neurons. Thus, MC3R in the BST may be acting antagonistically with CRH in response to stress and synergistically with GLP1R to decrease feeding. Future work will be required to distinguish between these possibilities.

Manipulations of BST circuitry have been shown to have robust effects on food intake and reward^3,18,19,54,55^. Optogenetic activation of AgRP fibers in the BST stimulates food intake in fed mice^55^, and antagonizing CRF2 receptors within the BST increases feeding following restraint stress^56^. Here, we found BST^MC3R^ neurons are activated to a greater degree in a fed state than a fasted state, and chemogenetic stimulation of BST^MC3R^ neurons decreases feeding in the fed state only in males. Our observation that BST^MC3R^ neurons are preferentially activated in a fed state, and exhibit the greatest increase in activity during the act of biting high fat chow, indicates BST^MC3R^ neurons may be preferentially involved in motivational and consummatory aspects of feeding. This corresponds well with previous data illustrating that optogenetic stimulation of BST GABAergic projections to the LHA results in a strong preference for high-fat food, even in a fed state^57^. Our finding that BST^MC3R^ neurons send dense projections to the LHA suggests that GABAergic BST^MC3R^ neurons may contribute to the reported high fat diet preference. Additionally, attraction to sweet stimuli encoded by the central amygdala is translated into consummatory responses in the BST^47^. Because BST^MC3R^ neurons receive dense afferents from the central amygdala, they may be involved in these consummatory responses.

In addition to being involved in feeding behaviors, BST^MC3R^ neurons also appear to be involved in responses to stress, including marked responses to looming stimuli, restraint, and shaker stress. The largest increase in BST^MC3R^ neuronal activity corresponding with any behavior occurred during responses to looming and shaking stress. Additionally, activation of BST^MC3R^ neurons significantly decreased struggling bouts during restraint stress in both males and females. An increased frequency of struggling during acute restraint is correlated with many other indices of stress and in humans is associated with a higher incidence of adverse health consequences, including gastric ulcers^45,58,59^. Therefore, struggling is typically classified as a coping response^35,45^, which implies that activating BST^MC3R^ neurons may dampen responses to restraint stress. A recent study reported AgRP neurons activate PVH^CRH^ neurons by inhibiting GABAergic BST afferents that typically restrain the activity of PVH^CRH^ neurons^60^. Since MC3R regulates AgRP signaling^32,61,62^, PVH^CRH^ neuronal activity is constrained by tonically active GABAergic afferents^63–66^, and BST^MC3R^ neurons are primarily GABAergic, projections to the PVH from BST^MC3R^ neurons may represent a broadly acting population of cells that restrain the activity of PVH^CRH^ neurons.

In contrast with the strong responses observed to stressors, we did not find evidence that BST^MC3R^ neurons are involved in anxiety-like behaviors. Because there are functionally distinct anxiogenic and anxiolytic neuronal populations in the dorsal BST^13,67^, it is possible that MC3R is expressed in both populations so as to cancel the effects of the other, which could explain why no effect was seen in any anxiety paradigm tested. Although intersectional approaches are required to parse out these effects, it is possible that repeated exposures to an anxiogenic environment are required before BST^MC3R^ neurons influence anxiety-like behaviors. In the present study, mice were only exposed to anxiogenic environments once. However, given the fact that other MC3R neuronal populations have been reported to modulate anxiety-like behavior^32,68^ in response to acute exposures, it is likely that BST^MC3R^ neurons may not play a defining role in driving generalized anxiety-like behaviors and instead are more crucial for modulating feeding and responses to stress.

In both chemogenetic and fiber photometry experiments, sexually dimorphic behavior was observed in association with feeding behavior, but not responses to stress. While BST^MC3R^ outputs were not sexually dimorphic, the density of afferents from several brain regions to BST^MC3R^ neurons were significantly different between males and females. Notably, most of the regions where males exhibited stronger inputs to BST^MC3R^ neurons reside in the thalamus and striatum and are known to regulate feeding^69–73^. Intriguingly, the parasubthalamic nucleus suppresses feeding^72,73^ and our data revealed that chemogenetic activation of BST^MC3R^ neurons decreased feeding in males that were in the fed state. Because BST^MC3R^ neurons in males received significantly stronger inputs from the parasubthalamic nucleus than that of females, it is possible that this input is influencing the decrease in feeding that was observed only in males. In contrast to males, most regions with stronger inputs to BST^MC3R^ neurons in females are located in the septum and hypothalamus. These regions contain high densities of estrogen receptors^74^ and are known to play a role in social behavior, aggression, and maternal behavior^75–82^. Altogether, denser afferents from regions associated with feeding, stress, motivation, and reward suggests BST^MC3R^ neurons in males may play a stronger role in integrating metabolic state with threat assessment and motivated behavior, whereas in females stronger afferents from regions associated with reproductive and social behaviors, indicates that BST^MC3R^ neurons may be more important for integrating metabolic state with reproductive status and social contexts.

Taken together, these data reveal a BST^MC3R^ neural circuit that is sexually dimorphic and impacts feeding, defensive behaviors and responses to stress. Although it remains unclear how each subpopulation of BST^MC3R^ neurons participates in context-specific regulation of downstream effector regions, these results may aid in explaining the gender disparity in the prevalence and treatment of stress- and eating-related disorders observed clinically, as well as provide a potential target for the treatment of these disorders.

## ACKNOWLEDGEMENTS

This work was supported by NIH grants F32DK123879 (MNB), K99/R00DK133560 (MNB), K99AA031509 (MAD), F32AA031404 (CME), R37AA019455 (DGW), P60AA031124 (DGW), R01DA042475 (DGW), and R01DK106476 (RBS).

## AUTHOR CONTRIBUTIONS

MNB and RBS initiated the project and prepared the manuscript with comments from all authors. MNB, RBS, DGW, MAD, and CME designed experiments. MNB, MAD, NP, and NTL analyzed experiments. MNB, MAD, CME, DS, SHH, HMR, JAB, and ASV performed experiments.

## COMPETING INTERESTS STATEMENT

The authors declare no competing interests.

## Notes

### Competing Interest Statement

The authors have declared no competing interest.

