## Supplemental Figures for "Sex-specific control of feeding and defensive behaviors by MC3R neurons in the bed nuclei of the stria terminalis"

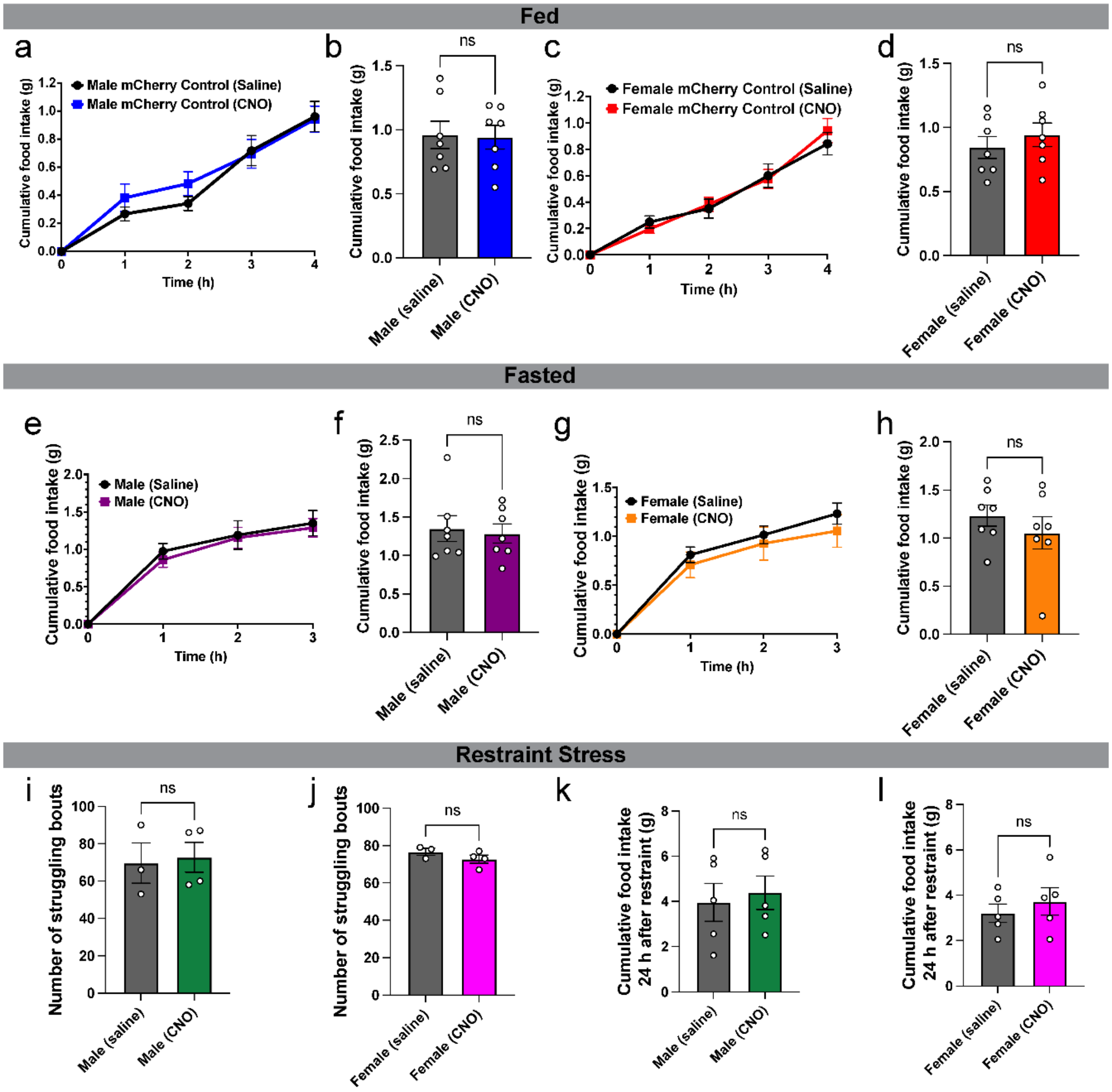


**Figure S1. mCherry DREADD control experiments.**

AAV5.mCherry was injected into the BSTd of male and female MC3R-Cre mice. Animals were allowed to recover for at least two weeks prior to any experiments being conducted. On the day of the experiment, mice received an injection of either CNO (2mg/kg) or saline 30 minutes prior to the beginning of the behavioral paradigm. **a-l** CNO treatment had no effect on **a-d** feeding in a fed state **e-h** feeding in a fasted state or **i-l** responses to restraint stress.
